# Epidemiologal investigation of an *Acinetobacter baumannii* outbreak using Core Genome Multilocus Sequence Typing

**DOI:** 10.1101/335307

**Authors:** Carolina Venditti, Antonella Vulcano, Silvia D’Arezzo, Cesare Ester Maria Gruber, Marina Selleri, Mario Antonini, Simone Lanini, Alessandra Marani, Vincenzo Puro, Carla Nisii, Antonino Di Caro

**Author notes:** **Corresponding author:** Vincenzo Puro, The National Institute for Infectious Diseases “L. Spallanzani”, Via Portuense 292, 00149 Rome, Italy.

## Abstract

Carbapenem-resistant (CR) *Acinetobacter baumannii* is a serious nosocomial pathogen able to cause a variety of serious, often life-threatening infections and outbreaks. We aimed to investigate the molecular epidemiology of clinical isolates of CR-*A. baumannii* from an outbreak occurred in the intensive care unit (ICU) of our hospital.

From January to April 2017, 13 CR-*A. baumannii* isolates were collected at the “L. Spallanzani” hospital, Rome, Italy; typing was performed by repetitive extragenetic palindromic-based (rep)- PCR-based DiversiLab system and WGS data were used for in *silico* analysis of traditional MLST types, for identifying resistance genes and for core genome multi locus sequence typing (cgMLST) analysis. Epidemiological data were obtained from hospital records.

Strains were cultured from 7 patients treated in the ICU of our hospital; all isolates showed a multi-drug resistant (MDR) profile, carrying the *bla*_OXA-23_ carbapenemase. Typing performed by rep-PCR and MLST showed that the isolates clustered into one group while the cgMLST approach, which uses 2690 gene targets to characterize the gene-by-gene allelic profile of *A. baumannii,* highlighted the presence of two cluster types. These results allowed us to identify two patients who entered the ICU already colonized by two different strains of CR-*A. baumannii*; we hypothesize that these two patients could be the source of two separate transmission chains.

Our results show that whole-genome-DNA sequencing by cgMLST is a valuable tool, better suited for prompt epidemiological investigations than traditional typing methods because of its higher discriminatory ability in determining clonal relatedness.

## Introduction

Adapted to persist in healthcare settings, *Acinetobacter baumannii* is one of the most successful pathogens in causing nosocomial infections, from respiratory diseases to bacteraemia and meningitis (1). It is often isolated in Intensive Care Units (ICUs) where its frequent resistance to carbapenems limits treatment options, increasing the risk of adverse outcomes for patients (2). The ability to survive in the environment and to acquire resistance determinants also allowed *A. baumannii* to become one of the most frequent causes of hospital outbreaks among Gram-negative pathogens (3, 4).

In recent years, several typing methods have been used to recognise distinct clonal lineages of *A. baumannii* during hospital outbreaks. The newer WGS-based methods, characterized by a higher discriminatory power, are now replacing standard approaches such as the traditional MLST and semi-automated repetitive extragenic palindromic-PCR (rep-PCR, DiversiLab) (5-7). Some studies carried out to investigate the epidemiology of Carbapenem-Resistant (CR)-*A. baumannii* outbreaks have shown that the WGS-based method known as core genome MLST (cgMLST) demonstrated good typing ability by extending the traditional MLST concept to the entire genome, thus offering additional information on genetic diversity (8-10). This method has been used for several bacteria causing outbreaks including, among others, *Klebsiella pneumoniae, Neisseria meningitis and Listeria monocytogenes* (11-13).

In this paper, we describe the use of the cgMLST technique to investigate an outbreak of CR-*A. baumannii* occurred in the intensive care unit (ICU) of our hospital.

## Methods

### Study design

Between December 2016 and May 2017, we analysed strains of CR-*A. baumannii* isolated from 7 critically ill patients treated in the ICU of the “L. Spallanzani” hospital in Rome, Italy, to investigate a potential clonal spread. According to the hospital’s surveillance programme aimed at detecting carbapenem-resistant bacteria, rectal swabs were taken from all patients upon admission, and then once weekly for the entire duration of their stay. The isolates included in this study were obtained from these surveillance samples as well as from infection sites. The strains were characterized using the typing methods described below, and their profiles were compared with those belonging to the Global Clone 1 and 2 (GC1 and GC2), which are dominant in Europe (14). Epidemiological data, including time of infection and transfer to other wards, movements of patients within the ICU and rooms occupied, were extracted from hospital records.

### Identification and Susceptibility testing

Samples were cultured on a selective medium designed to screen for carbapenemase production (chromID CARBA, bioMèrieux, Marcy l’Etoile, France); species identification and antimicrobial susceptibility were obtained by the Vitek-2 system (BioMérieux). As per EUCAST recommendations, MICs for tigecycline and colistin were confirmed by Etest (BioMérieux) and microbroth dilution methods (UMIC, Arnika Biocentric, Bandol, France), respectively (15).

### Rep-PCR

The UltraClean Microbial DNA Isolation Kit (Mo Bio Laboratories, Carlsbad, CA, USA) was used for genomic DNA extraction, and molecular typing by rep-PCR was performed as previously described; (16) results were analysed using the 2100 expert software. The presence of clusters was investigated by Pearson’s correlation coefficient and the unweighted pair-group method using average linkages (UPGMA). The guidelines followed for strain-level discrimination were those provided by the manufacturer: strains were considered as indistinguishable in case of a >97% similarity (no differences in fingerprints), as similar for a >95% similarity (1-2 band difference in fingerprints) and different if the similarity was <95%. The optimal cut-off chosen for cluster definition was 95% (6 Higg12). The Global Clones 1 and 2 were included in the analysis.

### WGS, assembly and cgMLST

DNA was quantified using the Qubit 2.0 fluorometer (Life Technologies, Carlsbad, CA, USA). Sequence data for all strains were obtained by the Illumina MiSeq System with 250-bp paried-end reads; DNA libraries were prepared using a Nextera XT DNA sample preparation kit (Illumina, San Diego, CA, USA), according to manufacturer’s instructions. Raw sequence data were submitted to the Sequence Read Archive (SRA) (Table 1). Sequence quality trimming was carried out as described by Bletz (17); d*e novo* assembly was performed by use of the Velvet assembler software (v1.1.04), integrated in the RIDOM SEQSPHERE+ software (version 2.1, Ridom GmbH, Münster, Germany).

**Table 1.** Summary of the clinical data and molecular characterization of CR-*A. baumannii* isolates.

| Patient | Date of admission | Date of isolation | Date of discharge from ICU | Strain | Ward | Sample | β-lactamase | Genes encoding non-β-lactam resistance | ST 'Pasteur' | ST 'Oxford' | DiversiLab Group-Type | cgMLST Cluster-Type | SRA Accession Number |
| --- | --- | --- | --- | --- | --- | --- | --- | --- | --- | --- | --- | --- | --- |
| P-1 * | 24/12/2016 | 24/12/2016 | 19/01/2017 | AC0* | ICU | Rectal swab | ND | ND | ND | ND | A | ND | ND |
|  |  | 10/01/2017 |  | AC1 | ICU | Bronchoaspirate | *bla*<sub>OXA-66</sub>, *bla*<sub>OXA-23</sub>, *bla*<sub>ADC-25</sub> | *aph(3')-VI-a*, *aph(6')-Id*, *armA*, *strA*, *mph(E)*, *mrs(E)*, *sul2*, *tet(B)* | 2 | 451 | A | 1 | SRR6436005 |
| P-2 | 20/01/2017 | 02/02/2017 | 13/02/2017 | AC2 | ICU | Rectal swab | *bla*<sub>OXA-66</sub>, *bla*<sub>OXA-23</sub>, *bla*<sub>ADC-25</sub> | *aph(3')-VI-a*, *aph(6')-Id*, *armA*, *strA*, *mph(E)*, *mrs(E)*, *sul2*, *tet(B)* | 2 | 451 | A | 1 | SRR6436006 |
|  |  | 09/02/2017 |  | AC3 | ICU | Blood | *bla*<sub>OXA-66</sub>, *bla*<sub>OXA-23</sub>, *bla*<sub>ADC-25</sub> | *aph(3')-VI-a*, *aph(6')-Id*, *armA*, *strA*, *mph(E)*, *mrs(E)*, *sul2*, *tet(B)* | 2 | 451 | A | 1 | SRR6436007 |
| P-3 | 10/02/2017 | 25/02/2017 | 07/03/2017 | AC4 | ICU | Blood | *bla*<sub>OXA-66</sub>, *bla*<sub>OXA-23</sub>, *bla*<sub>ADC-25</sub> | *aph(3')-VI-a*, *aph(6')-Id*, *armA*, *strA*, *mph(E)*, *mrs(E)*, *sul2*, *tet(B)* | 2 | 451 | A | 1 | SRR6436008 |
|  |  | 15/03/2017 |  | AC10 | CW | Bronchoaspirate | *bla*<sub>OXA-66</sub>, *bla*<sub>OXA-23</sub>, *bla*<sub>ADC-25</sub> | *aph(3')-VI-a*, *aph(6')-Id*, *armA*, *strA*, *mph(E)*, *mrs(E)*, *sul2*, *tet(B)* | 2 | 451 | A | 1 | SRR6436000 |
| P-4 * | 11/03/2017 | 11/03/2017 | 09/04/2017 | AC7 | ICU | Rectal swab | *bla*<sub>OXA-66</sub>, *bla*<sub>OXA-23</sub>, *bla*<sub>ADC-25</sub> | *aph(3')-VI-a*, *aph(6')-Id*, *armA*, *strA*, *mph(E)*, *mrs(E)*, *sul2*, *tet(B)* | 2 | 451 | A | 2 | SRR6436003 |
|  |  | 13/03/2017 |  | AC8 | ICU | Bronchoaspirate | *bla*<sub>OXA-66</sub>, *bla*<sub>OXA-23</sub>, *bla*<sub>ADC-25</sub> | *aph(3')-VI-a*, *aph(6')-Id*, *armA*, *strA*, *mph(E)*, *mrs(E)*, *sul2*, *tet(B)* | 2 | 451 | A | 2 | SRR6436004 |
| P-5 | 05/03/2017 | 16/03/2017 | 21/03/2017 | AC9 | ICU | Rectal swab | *bla*<sub>OXA-66</sub>, *bla*<sub>OXA-23</sub>, *bla*<sub>ADC-25</sub> | *aph(3')-VI-a*, *aph(6')-Id*, *armA*, *strA*, *mph(E)*, *mrs(E)*, *sul2*, *tet(B)* | 2 | 451 | A | 1 | SRR6435999 |
|  |  | 24/03/2017 |  | AC11 | CW | Blood | *bla*<sub>OXA-66</sub>, *bla*<sub>OXA-23</sub>, *bla*<sub>ADC-25</sub> | *aph(3')-VI-a*, *aph(6')-Id*, *armA*, *strA*, *mph(E)*, *mrs(E)*, *sul2*, *tet(B)* | 2 | 451 | A | 1 | SRR6435994 |
| P-6 | 16/03/2017 | 23/03/2017 | 27/03/2017 | AC12 | ICU | Rectal swab | *bla*<sub>OXA-66</sub>, *bla*<sub>OXA-23</sub>, *bla*<sub>ADC-25</sub> | *aph(3')-VI-a*, *aph(6')-Id*, *armA*, *strA*, *mph(E)*, *mrs(E)*, *sul2*, *tet(B)* | 2 | 451 | A | 1 | SRR6435995 |
|  |  | 28/03/2017 |  | AC13 | CW | Blood | *bla*<sub>OXA-66</sub>, *bla*<sub>OXA-23</sub>, *bla*<sub>ADC-25</sub> | *aph(3')-VI-a*, *aph(6')-Id*, *armA*, *strA*, *mph(E)*, *mrs(E)*, *sul2*, *tet(B)* | 2 | 451 | A | 1 | SRR6435996 |
| P-7 | 10/04/2017 | 20/04/2017 | 11/05/2017 | AC15 | ICU | Bronchoaspirate | *bla*<sub>OXA-66</sub>, *bla*<sub>OXA-23</sub>, *bla*<sub>ADC-25</sub> | *aph(3')-VI-a*, *aph(6')-Id*, *armA*, *strA*, *mph(E)*, *mrs(E)*, *sul2*, *tet(B)* | 2 | 451 | A | 2 | SRR6435997 |
|  |  | 30/04/2017 |  | AC16 | ICU | Blood | *bla*<sub>OXA-66</sub>, *bla*<sub>OXA-23</sub>, *bla*<sub>ADC-25</sub> | *aph(3')-VI-a*, *aph(6')-Id*, *armA*, *strA*, *mph(E)*, *mrs(E)*, *sul2*, *tet(B)* | 2 | 451 | A | 2 | SRR6435998 |

\* Patients colonised prior to admission.
\* Molecular characterization and typing was not performed on rectal swab Patient 1 (study code AC0).
Abbreviation: ICU, Intensive Care Unit; CW, Clinical ward; ND, not done; SRA, Sequence Read Archive.

In-depth phylogenetic analysis was achieved by cgMLST, using the software with default parameters (17). To generate an *ad hoc* scheme, Open Reading Frames (ORFs) in the genome of each isolate were identified by SeqSphere, using the whole-genome AB042 *A. baumannii* strain as reference (GenBank Accession number CP019034.1). Briefly, 2690 targets genes were used to characterize the gene-by-gene allelic profile of the *A. baumannii* strains, under the assumption that a well-defined cgMLST scheme should cover at least 95% of the genes present in all analysed isolates, compared to the reference strain. The resulting set of target genes was then used for interpreting the clonal relationship displayed in a minimum spanning tree (MST) using the “pairwise ignore missing values” parameter during distance calculations.

In accordance with other studies, we decided to set the cluster definition threshold at a maximum difference of 12 alleles in a pairwise comparison (8, 9).

### MLST and resistance genes identification

The identified ORF were uploaded to the ResFinder v3.0 web server for identifying resistance genes and the MLST web server for the Pasteur and Oxford scheme MLST analysis (www.genomicepidemiology.org) (18).

## Results and discussion

Of the 13 isolates studied, nine were cultured from clinically relevant samples, and the remaining four were obtained from rectal swabs. Only the first two isolates from each patient were included in the analysis, with one exception (Patient 1) for whom only one strain (isolated from a bronchoaspirate 17 days after admission) was available. For patients 3, 5 and 6, the second isolate was collected once the patient had been transferred from the ICU to a clinical ward. Two of the 7 patients, (1 and 4) were colonised prior to ICU admission, as determined by their positive rectal swabs (Table 1).

All strains were resistant to amoxicillin/clavulanic acid (MIC ≥ 32 mg/L), cefotaxime (MIC ≥ 64 mg/L), ciprofloxacin (MIC ≥ 4 mg/L), ertapenem (MIC ≥ 8 mg/L), imipenem (MIC ≥ 16 mg/L), gentamicin (MIC ≥ 16 mg/L) and trimethoprim/sulfamethoxazole (MIC ≥ 320 mg/L); colistin MICs ranged from 0.5-0.75 mg/L (all strains were susceptible), while the MICs for tigecycline were between 0.5 and 1.5 mg/L. Also the resistance profile obtained from the i*n silico* ResFinder analysis showed identical resistance gene profiles for all isolates, as well as the presence of the following genes: *bla*_OXA-66_ β-lactamase (an OXA enzyme belonging to the intrinsic OXA-51-like enzymes) (19), *bla*_OXA-23_ carbapenemase, and the cephalosporinase-encoding *bla*_ADC-25_ (Table 1). In addition, all isolates were shown to belong to the Sequence Type (ST)-2 and ST451using the Pasteur and Oxford scheme respectively. Typing performed by rep-PCR produced a similar result; all clinical isolates clustered into one group (Group-A) displaying a > 95% similarity with GC2, and only 70% with GC1, as expected since GC2 is the most prevalent clone in Europe and has been involved in the majority of outbreaks (20) (see Fig. S1 in the supplemental material).

For a more in-depth analysis, genotyping results obtained by MLST schemes and rep-PCR were compared with those of the cgMLST method. Sequences were obtained with a median mean coverage of 108 (range: 67-115) for all strains. The final cgMLST scheme consisted of 2690 target genes; of these, 2499 were consistently present. The percentage of good targets, based on the core genome, ranged between 96.9% and 98.4% (median 98.3%). These results show the presence of two clusters types (CT-1 and CT-2) differentiated by only 17 allelic differences, where the traditional methods described above had highlighted only one (Figure 1). Nine isolates grouped within CT-1, (AC1 from P-1, AC2 and AC3 from P-2, AC4 and AC5 from P-3, AC8 and AC9 from P-5 and AC10 and AC11 from P-6), while 4 strains isolated from 2 patients (P-4 and P-7) gave rise to CT-2 (AC6, AC7, AC12 and AC13). The strains within each cluster showed a very high level of correlation; both clusters displayed a good similarity with GC2 (up to 252 differences), and only a distant correlation with GC1 (>2000 differences) (Figure 1).

**Figure 1.**
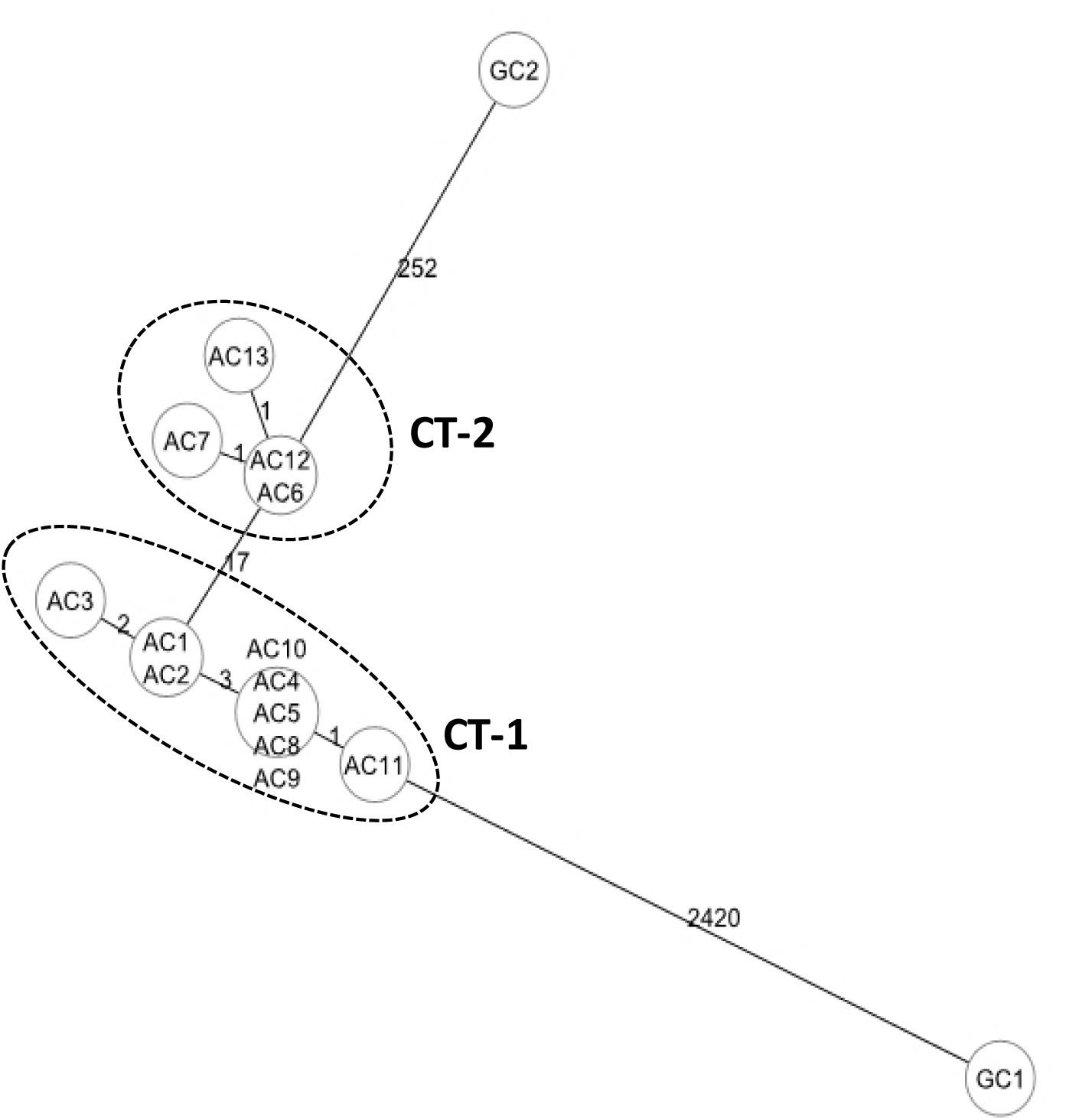
Minimum spanning tree based on cgMLST analysis of 13 CR-*A*. *baumannii* isolates (AC1 to AC13) and Global Clones 1 and 2 strains (GC1 and GC2). Each circle represents an allelic profile, i.e. genotype, based on sequence analysis of up to 2,690 target genes. The numbers on the connecting lines illustrate the numbers of target genes with different alleles. Cluster types (grouped together in the dotted ellipses) consist of closely related genotypes (≤12 allele differences) and are numbered consecutively (I and II).

The presence in each cluster of a patient colonised prior to admission (patients 1 and 4), and the fact that the same clones were later found in other patients treated in the ICU, suggested that two distinct transmission events occurred. To support this hypothesis, hospital records were studied to determine how the patients were treated and moved within the ICU. The ward is composed of nine single rooms; the 7 patients occupied rooms 2, 3, 4, and 5, which are next to each other. Figure 2 shows the timeline of patients and room occupancy during the five-month study period, and the days in which the patients were negative or positive for CR-*A. baumannii*. For cluster-type 1, we hypothesize a chain transmission involving patients 1, 2, 3, 5, and 6, as suggested by the brief overlap between the end of one patient’s stay and the beginning of the next (Figure 2). Cluster-type 2 could have originated from a separate transmission from patient 4 to patient 7, both hosted in room 5.

**Figure 2.**
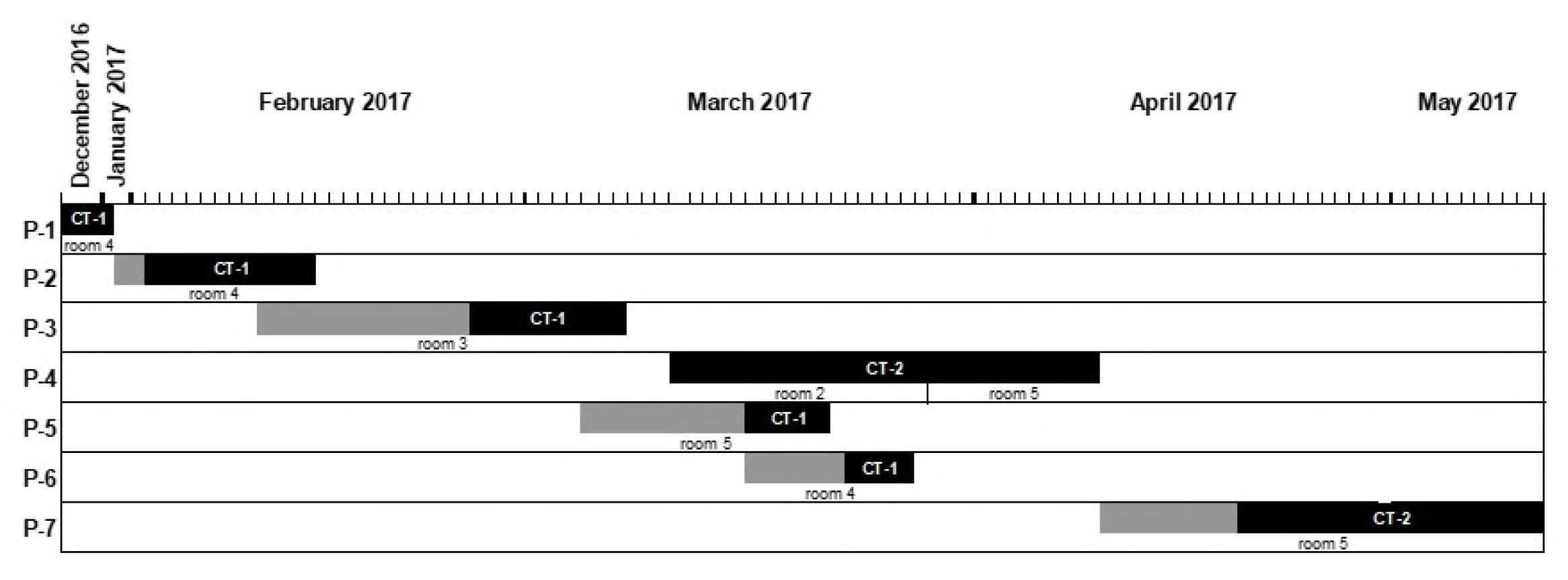
Timeline and room occupancy of all 7 patients (P-1 to P-7) positive for CR-*A*. *baumannii* during December 2016 and May 2017. All patients occupied the same room for the entire duration of their stay, except patient 4 who was moved from room 2 to 5 after 18 days. Grey boxes: days in which patients were negative for CR-*A*. *baumannii* Black boxes: days in which patients were positive for CR-*A*. *baumannii* CT-1: Cluster type 1 CT-2: Cluster type 2

Our study did not include environmental samples, which could have been useful in identifying a possible role of the environment in the transmission chain (21). However the genotyping results and the epidemiological data support the hypothesis that the origin of the two distinct clusters could be traced back to patients 1 and 4, who had been admitted to the ICU with rectal swabs already positive for CR-*A. baumannii*.

Investigations were undertaken to highlight possible breaches in the isolation precautions employed, but nothing was found that justified any changes in the hospital’s infection control procedures.

Every effort was made to contain the outbreak and our continued surveillance showed, to this day, no further cases either in the ICU or in the clinical wards where the patients were later transferred. In conclusion, our data are in agreement with other recent reports (8, 9) that highlight how WGS-based methods such as the cgMLST represent reliable techniques that will likely become the gold standard for strain subtyping in support of epidemiological investigations.

## Supplemental material

**Figure S1.** Dendrogram generated by the DiversiLab software of rep-PCR-banding patterns for the13 CR-*A. baumannii* isolates (AC1 to AC13) and Global Clones 1 and 2 strains (GC1 and GC2) included for comparison. Using a similarity threshold of 95% to define a cluster, the resulting rep-PCR cluster is named A.

Raw sequence data were submitted on Sequence Read Archive (SRA) under the BioProject PRJNA427128, SRA Study: SRP127890, Accession numbers Sequences see Table1.

## Funding Information

This work was supported by “Ricerca corrente” and “5×1.000” research funds from the Italian Ministry of Health.

## Transparency declarations

None of the authors declared any conflicts of interest.

## References

1. Munoz-Price LS, Weinstein RA. 2008. Acinetobacter infection. N Engl J Med. 20;358(12):1271–81. DOI:10.1056/NEJMra070741.

2. Peleg AY, de Breij A, Adams MD, Cerqueira GM, Mocali S, Galardini M, Nibbering PH, Earl AM, Ward DV, Paterson DL, Seifert H, Dijkshoorn L. 2012. The success of acinetobacter species; genetic, metabolic and virulence attributes. PLoS One. 7(10):e46984. DOI:10.1371/journal.pone.0046984.

3. Jawad A, Seifert H, Snelling AM, Heritage J, Hawkey PM. 1998. Survival of Acinetobacter baumannii on dry surfaces: comparison of outbreak and sporadic isolates. J Clin Microbiol. Jul;36(7):1938–41.

4. Husni RN, Goldstein LS, Arroliga AC, Hall GS, Fatica C, Stoller JK, Gordon SM. 1999. Risk factors for an outbreak of multi-drug-resistant Acinetobacter nosocomial pneumonia among intubated patients. Chest. 115(5):1378–82.

5. Zarrilli R, Pournaras S, Giannouli M, Tsakris A. 2013. Global evolution of multidrug-resistant Acinetobacter baumannii clonal lineages. Int J Antimicrob Agents. 41(1):11–9. DOI:10.1016/j.ijantimicag.2012.09.008.

6. Higgins PG, Hujer AM, Hujer KM, Bonomo RA, Seifert H. 2012. Interlaboratory reproducibility of DiversiLab rep-PCR typing and clustering of Acinetobacter baumannii isolates. J Med Microbiol. 61(Pt 1):137–41. DOI:10.1099/jmm.0.036046-0.

7. Hammerum AM, Hansen F, Skov MN, Stegger M, Andersen PS, Holm A, Jakobsen L, Justesen US. 2015. Investigation of a possible outbreak of carbapenem-resistant Acinetobacter baumannii in Odense, Denmark using PFGE, MLST and whole-genome-based SNPs. J Antimicrob Chemother. 70(7):1965–8. DOI:10.1093/jac/dkv072.

8. Willems S, Kampmeier S, Bletz S, Kossow A, Köck R, Kipp F, Mellmann A. 2016. Whole-Genome Sequencing Elucidates Epidemiology of Nosocomial Clusters of Acinetobacter baumannii. J Clin Microbiol. 54(9):2391–4. DOI:10.1128/JCM.00721-16.

9. Higgins PG, Prior K, Harmsen D, Seifert H. 2017. Development and evaluation of a core genome multilocus typing scheme for whole-genome sequence-based typing of Acinetobacter baumannii. PLoS One. 8;12(6):e0179228. DOI:10.1371/journal.pone.0179228.

10. Dekker JP, Frank KM. 2016. Next-Generation Epidemiology: Using Real-Time Core Genome Multilocus Sequence Typing To Support Infection Control Policy. J Clin Microbiol.54(12):2850–2853.

11. Bialek-Davenet S, Criscuolo A, Ailloud F, Passet V, Jones L, elannoy-Vieillard AS, Garin B, Le Hello S, Arlet G, Nicolas-Chanoine MH, Decré, Brisse S. 2014. Genomic definition of hypervirulent and multidrug-resistant lebsiella pneumoniae clonal groups. Emerg Infect Dis. 20(11):1812–20. DOI:10.3201/eid2011.140206.

12. Bratcher HB, Corton C, Jolley KA, Parkhill J, Maiden MC. 2014. A gene-by-gene population genomics platform: de novo assembly, annotation and genealogical analysis of 108 representative Neisseria meningitidis genomes. BMC Genomics. 18;15:1138. DOI:10.1186/1471-2164-15-1138.

13. Ruppitsch W, Pietzka A, Prior K, Bletz S, Fernandez HL, Allerberger F, Harmsen D, Mellmann A. 2015. Defining and Evaluating a Core Genome Multilocus Sequence Typing Scheme for Whole-Genome Sequence-Based Typing of Listeria monocytogenes. J Clin Microbiol. 53(9):2869–76. DOI:10.1128/JCM.01193-15.

14. Mak JK, Kim MJ, Pham J, Tapsall J, White PA. 2009. Antibiotic resistance determinants in nosocomial strains of multidrug-resistant Acinetobacter baumannii. J Antimicrob Chemother. 63(1):47–54. DOI:10.1093/jac/dkn454.

15. European Committee on Antimicrobial Susceptibility Testing (EUCAST). Breakpoint tables for interpretation of MICs and zone diameters. Version 8.0 EUCAST; 2018. Available from: http://www.eucast.org/clinical-breakpoints/

16. Fitzpatrick MA, Ozer EA, Hauser AR. 2016. Utility of Whole-Genome Sequencing in Characterizing Acinetobacter Epidemiology and Analyzing Hospital Outbreaks. J Clin Microbiol. 54(3):593–612. DOI:10.1128/JCM.01818-15.

17. Bletz S, Mellmann A, Rothgänger J, Harmsen D. 2015. Ensuring backwards compatibility: traditional genotyping efforts in the era of whole genome sequencing. Clin Microbiol Infect. 21(4):347.e1-4. DOI:10.1016/j.cmi.2014.11.005.

18. Larsen MV, Cosentino S, Rasmussen S, Friis C, Hasman H, Marvig RL, Jelsbak L, Sicheritz-Pontén T, Ussery DW, Aarestrup FM, Lund O. 2012. Multilocus sequence typing of total-genome-sequenced bacteria. J Clin Microbiol. 50(4):1355–61. DOI:10.1128/JCM.06094-11.

19. Woodford N, Ellington MJ, Coelho JM, Turton JF, Ward ME, Brown S, Amyes SG, Livermore DM. 2006. Multiplex PCR for genes encoding prevalent OXA carbapenemases in Acinetobacter spp. Int J Antimicrob Agents. 27(4):351–3.

20. Tomaschek F, Higgins PG, Stefanik D, Wisplinghoff H, Seifert H. 2016. Head-to-Head Comparison of Two Multi-Locus Sequence Typing (MLST) Schemes for Characterization of Acinetobacter baumannii Outbreak and Sporadic Isolates. PLoS One. 12;11(4):e0153014. DOI:10.1371/journal.pone.0153014.

21. Chemaly RF, Simmons S, Dale C Jr, Ghantoji SS, Rodriguez M, Gubb J, Stachowiak J, Stibich M. 2014. The role of the healthcare environment in the spread of multidrug-resistant organisms: update on current best practices for containment. Ther Adv Infect Dis. 2(3-4):79–90. DOI:10.1177/2049936114543287.

